# Disturbed proteostasis is the key event that triggers cellular senescence through the regulation of chromatin organization and genome stability

**DOI:** 10.1101/2025.03.03.641349

**Authors:** Yuki Takauji, Keisuke Tanabe, Haruna Miyashita, Takuma Katakawa, Saeri Fujii, Atsuki En, Michihiko Fujii

## Abstract

Mammalian cells undergo irreversible proliferation arrest when exposed to stresses, a phenomenon termed cellular senescence. Various types of stress induce cellular senescence; nonetheless, senescent cells show similar phenotypes overall. Thus, cells undergo cellular senescence through the similar mechanisms, regardless of the type of stress encountered. Here we aimed to reveal the mechanisms underlying cellular senescence. We have previously shown that lamin b receptor (LBR), which is a protein that regulates heterochromatin organization, was downregulated in senescent cells, and downregulation of LBR induced cellular senescence. Additionally, we have shown that downregulation of protein synthesis effectively suppressed cellular senescence. Thereby, chromatin organization and protein synthesis are implicated in the regulation of cellular senescence. We examined the roles of them in cellular senescence and found that protein synthesis was upregulated during the induction of cellular senescence, and upregulated protein synthesis caused disturbed proteostasis that led to the decreased function of LBR. Furthermore, we showed that decreased LBR function induced cellular senescence through altered chromatin organization and increased genome instability. Importantly, these findings revealed a link between protein synthesis and chromatin organization and accounted for the phenotypes of senescent cells which show disturbed proteostasis, altered chromatin organization, and increased genome instability. Our findings provided the general model for the mechanisms of cellular senescence.

## 1 Introduction

Normal types of somatic cell undergo terminal proliferation arrest after a limited number of cell divisions, a phenomenon termed cellular senescence (Hayflick & Moorehead, 1961).

Cellular senescence is also induced by various types of stress such as DNA damage, oxidative stress, and activation of oncogenes in cells, regardless of normal or cancerous cell types (Campisi & d’Adda di Fagagna, 2007). Thus, cellular senescence is regarded as a phenomenon of irreversible cellular proliferation arrest when cells are exposed to stresses. Senescent cells increase in aged organisms, and elimination of senescent cells contributes to improving organismal health with extension of lifespans (Baker et al., 2016). Thus, prevention of the increase of senescent cells is beneficial to organismal health, and understanding of the mechanisms for cellular senescence is of importance.

Cellular senescence is induced by a broad range of stresses in various types of cell; nonetheless, senescent cells show similar phenotypes overall. This suggests that cells undergo cellular senescence through the similar mechanisms after being exposed to stresses, regardless of the type of cell or the type of stress. Senescent cells generally exhibit an enlarged and flattened cell morphology with upregulation of the senescence-associated β-galactosidase (SA-β-gal) activity and show altered gene expression, as represented by the upregulated expression of secretory factors (senescence-associated secretory phenotype, SASP) (Campisi & d’Adda di Fagagna, 2007). Senescent cells also show decreased maintenance of protein homeostasis (proteostasis) and genome stability (Chen, Hales, & Ozanne, 2007; Sabath et al., 2020). Additionally, altered organization of heterochromatin is a well-known feature of senescent cells (Narita et al., 2003; Sadaie et al., 2013; Swanson, Manning, Zhang, & Lawrence, 2013). This phenomenon is of particular interest, given that chromatin organization is involved in the regulation of fundamental genome functions such as gene expression, DNA replication, and maintenance of genome stability. Thus, altered chromatin organization is presumed to be involved in determining the characteristics of senescent cells.

Heterochromatin organization is regulated by the nuclear lamina proteins, lamin A/C and lamin B, and the nuclear envelope protein such as lamin B receptor (LBR). Notably, these proteins are shown to play important roles in cellular senescence. For instance, certain types of mutation in lamin A/C are responsible for the Hutchinson-Gilford progeria syndrome, and lamin B is downregulated in senescent cells (De Sandre-Giovannoli et al., 2003; Eriksson et al., 2003; Shimi et al., 2011). LBR is also downregulated in senescent cells, and indeed, *LBR* is determined to be one of the 18 RNAs that are most consistently downregulated in cellular senescence induced by various means in different cell types (Arai et al., 2019; Arai et al., 2016; Casella et al., 2019; Lukasova, Kovar ik, Bac ikova, Falk, & Kozubek, 2017). Furthermore, knockdown of *LBR* induces cellular senescence, and conversely, enforced expression of *LBR* suppresses it (Arai et al., 2019; En, Takauji, Miki, Ayusawa, & Fujii, 2020; Herman et al., 2021). *LBR* regulates SASP and genome stability, and its upregulated expression is involved in cellular immortalization (En, Takauji, Ayusawa, & Fujii, 2020; En, Takemoto, Yamakami, Nakabayashi, & Fujii, 2024). These findings suggest that LBR plays crucial roles in cellular senescence through the regulation of chromatin organization. However, the mechanisms by which chromatin organization regulates cellular senescence are not well determined.

In addition to chromatin organization, protein synthesis is involved in the regulation of cellular senescence. In the previous studies, we have shown that a low dose of cycloheximide, a protein synthesis inhibitor, effectively suppressed cellular senescence induced by various types of stress in both normal and cancerous cells (En, Takauji, Miki, et al., 2020; Sumikawa, Matsumoto, Sakemura, Fujii, & Ayusawa, 2005; Takauji, En, Miki, Ayusawa, & Fujii, 2016; Takauji, Wada, et al., 2016). Additionally, a low dose of cycloheximide extends the organismal lifespan of *Caenorhabditis elegans* (Takauji, Wada, et al., 2016). Since a low dose of cycloheximide slows down protein synthesis, these results suggest that the rate of protein synthesis may be one of the determinants of cellular senescence. However, the mechanisms by which protein synthesis regulates cellular senescence are not well determined.

Here we aimed to reveal the mechanisms underlying cellular senescence by analyzing the roles of chromatin organization and protein synthesis in cellular senescence. For this, human cells were induced to undergo cellular senescence by several means: *i.e.*, by serial passage (replicative senescence), by treatment with genotoxic stress (chemotherapy-induced senescence), or by expression of oncogenic Ras (oncogene-induced senescence). We showed that protein synthesis was upregulated during the induction of cellular senescence, and upregulated protein synthesis caused disturbed proteostasis that led to the decreased function of LBR. Then, decreased LBR function induced cellular senescence through altered chromatin configuration and increased genome instability. These results accounted for the phenotypes of senescent cells which show disturbed proteostasis, altered chromatin organization, and increased genome instability, and provided the general model for the mechanisms of cellular senescence.

## 2 Materials and Methods

### 2.1 Cell culture

Normal primary human fibroblast TIG-7 cells and human cervical cancer HeLa cells were purchased from the Japanese Collection of Research Bioresources (JCRB). Human SVts7-1 cells were generously provided by Prof. Ide, T., as previously described (Fujii et al., 1995). HeLa cells and SVts7-1 cells were cultured in DMEM medium (Nissui) supplemented with 5% fetal bovine serum (Sigma-Aldrich) on tissue culture dishes (Thermo Fisher Scientific) under 5% CO_2_ and 95% humidity. Similarly, TIG-7 cells were cultured in DMEM medium supplemented with 10% fetal bovine serum, and HeLa^T-LBR^ cells, a HeLa cell line that expresses *LBR* in a doxycycline-dependent manner, were cultured in DMEM medium supplemented with 7% fetal bovine serum and 0.4% glucose (En, Takauji, Miki, et al., 2020).

### 2.2 Cell proliferation assay

To determine the proliferative potential of cells, appropriate numbers of cells were plated on 35-mm dishes, and grown for 1-2 weeks. The colonies were visualized by staining with Coomassie Brilliant Blue (CBB, Bio-Rad).

### 2.3 DNA transfection

The plasmids that express Ras^V12^ or LBR were constructed by inserting their cDNAs in pcDNA3.1 (Invitrogen) (En, Takauji, Miki, et al., 2020). The plasmids were introduced into human cells by electroporation with a high-efficiency electroporator (type NEPA21, Nepa Gene). Cells (10^6^ cells) were transfected with the test plasmids (3-5 μg) together with the plasmid carrying the puromycin-resistance gene, and transfected cells were enriched by selection with puromycin for 2-4 days for analyses.

### 2.4 Cellular senescence

Cellular senescence was induced by treatment with camptothecin (17.5 nM) (Sigma-Aldrich) in HeLa cells, by serial passage in TIG-7 cells, or by transfection of the plasmid expressing oncogenic Ras^V12^ in SVts7-1 cells. To suppress cellular senescence, 4-phenyl butyric acid (4-PBA; Wako) was added to the cells at the optimal concentration for each cell type in each experiment. HeLa cells were treated with 300 μM or 30 μM (low dose) of 4-PBA; SVts7-1 cells were treated with 100 μM of 4-PBA; and TIG-7 cells were treated with 3-30 μM of 4-PBA. Similarly, cycloheximide (Wako) was added at 0.15 μM to HeLa cells.

### 2.5 Senescence-associated ß-Galactosidase assay

Cells were fixed with a fixation solution (2% formaldehyde and 0.2% glutaraldehyde) at room temperature for 5 min, and incubated with a staining solution (40 mM citric acid-sodium phosphate (pH 6.0), 150 mM NaCl, 2 mM MgC1_2_, 1 mg/ml of 5-bromo-4-chloro-3-indolyl ß-D-galactoside, 5 mM potassium ferricyanide, and 5 mM potassium ferrocyanide) at 37°C. After washing with PBS, the cells were photographed under a microscope.

### 2.6 Protein synthesis

Protein synthesis was measured by the Click-iT HPG Alexa Fluor Protein Synthesis Assay Kit according to the manufacturer’s instructions (Molecular probes). In brief, cells were cultured with L-homopropargylglycine (HPG), a methionine analogue, in methionine-free DMEM medium for 30 min to label nascent polypeptides. Cells were fixed with 3.7 % formaldehyde for 20 min, permeabilized with 0.5% Triton X-100 for 15 min, and processed with the Click-iT reaction cocktail for 30 min. Labelled polypeptides were visualized by fluorescence microcopy (Keyence, BZ-X710) and quantified with supplied software.

### 2.7 Detergent insoluble proteins

To isolate detergent-insoluble proteins, cells were collected in the RIPA buffer (20 mM Tris-HCl, 150 mM NaCl, 1% Nonidet P-40, 0.5% sodium deoxycholate, 0.1% SDS, 1 mM phenylmethanesulfonyl fluoride, 2 μg/ml leupeptin, 2 μg/ml aprotinin, 10 mM dithiothreitol), and disrupted by sonication for 10-15 sec on ice. The protein concentrations were determined (Takara), and the cell lysates containing equal amounts of proteins were centrifugated at 15 krpm for 10 min to collect the supernatants and the pellets. The pellets, after washing with the RIPA buffer two times, were recovered as detergent-insoluble proteins. These proteins were subjected to SDS-PAGE analysis and detected with SYPRO Ruby protein gel stain (Thermo Fisher Scientific).

### 2.8 ProteoStat

Protein aggregates were detected by the ProteoStat Aggresome Detection Kit according to the manufacturer’s instructions (Enzo Life Sciences). In brief, cells were fixed with 4% formaldehyde for 30 min, and permeabilized with 0.5% Triton X-100 for 30 min, and stained with the ProteoStat aggresome detection reagent for 30 min. Fluorescence signals were detected by fluorescence microscopy (Keyence, BZ-X710) and quantified with supplied software.

### 2.9 Indirect immunofluorescence analysis

Cells were cultured on a cover slip and fixed with methanol for 15 min at -20°C. The cells were incubated with bovine serum albumin (1%) at room temperature for 1 h and incubated with the primary antibody against LBR (Cosmo Bio) or γ-H2AX (Cell Signaling) for 16-24 h. Subsequently, the cells were incubated with an alexa 568-conjugated or alexa 546-conjugated secondary antibody (Molecular Probes) for 3 h and with DAPI (4’,6-diamidino-2-phenylindole, Wako) for 30 min, and mounted with an anti-fading reagent (Molecular probes). Fluorescence images were captured by fluorescence microscopy (BZ-X710, Keyence) and quantified with supplied software.

### 2.10 DNase I sensitivity assay

Nuclease sensitivity assays were performed as previously described (En et al., 2024; En, Watanabe, Ayusawa, & Fujii, 2022). Cells were permeabilized for 5 min at room temperature with the buffer [10 mM PIPES (pH 7.0), 100 mM NaCl, 300 mM sucrose, 3 mM MgCl_2_, and 0.2% Triton X-100], and then treated with DNase I (5-10 U/ml) for 20 min at 37℃ in the buffer [10 mM Tris-HCl (pH7.5), 0.5 mM CaCl_2_, 2.5 mM MgCl_2_, 10 mM PIPES (pH7.0), 100 mM NaCl, and 300 mM sucrose]. These cells were fixed with formaldehyde and stained with DAPI. The fluorescence signals of DAPI were detected by fluorescence microscopy (BZ-X710, Keyence) and quantified with supplied software.

### 2.11 Statistical analysis

Data are presented as mean ± standard deviation. Statistical analysis was performed with SPSS software (IBM).

### 2.12 Quantitative RT-PCR

RNA was prepared from cells by the acid guanidinium thiocyanate-phenol-chloroform extraction method, and cDNA was synthesized by a reverse transcriptase (PrimeScript 1st strand cDNA synthesis kit, Takara). The transcript of the gene was quantified by quantitative real time PCR with a kit (Thunderbird SYBR qPCR Mix, Toyobo) on a StepOnePlus Real-Time PCR System (Applied Biosystems) according to the supplier’s manuals. The primers used were as follows: 5’-CAGAACCAGCAGAGGTCACA-3’ and 5’-AGCTGTGCCACTTTCCTTTC-3’ for *CHOP*; 5’-TGCTGAGTCCGCAGCAGGTG -3’ and 5’-GCTGGCAGGCTCTGGGGAAG-3’ for *XBP-1*; 5’-CTGGGTACATTTGATCTGACTGG-3’ and 5’-GCATCCTGGTGGCTTTCCAGCCAT TC-3’ for *GRP78*; 5’-GAAGGTGAAGGTCGGAGTCAA-3’ and 5’-GACAAGCTTCCCGTTCTCAG-3’ for *GAPDH.* The expression level of each gene was normalized with the expression of *GAPDH*.

## 3 Results

### 3.1 Upregulated protein synthesis during the induction of cellular senescence

We have previously shown that a low dose of cycloheximide, which slows down protein synthesis, effectively suppressed the induction of cellular senescence in human primary fibroblast TIG-7 cells and HeLa cells (En, Takauji, Miki, et al., 2020; Sumikawa et al., 2005; Takauji, En, et al., 2016; Takauji, Wada, et al., 2016). This finding suggested a role of protein synthesis in cellular senescence; however, the mechanisms by which protein synthesis regulates cellular senescence were not well identified. We first examined whether protein synthesis is altered upon induction of cellular senescence. Cellular senescence was induced by treatment of HeLa cells with camptothecin (CPT), a DNA topoisomerase inhibitor that causes genotoxic stress by production of DNA strand breaks (Fig. 1A). We found that protein synthesis was upregulated during the induction of cellular senescence by CPT (Figs. 1B and C); and that slowdown of protein synthesis by a low dose of cycloheximide, which did not largely affect cell proliferation, suppressed the upregulation of protein synthesis as well as the induction of cellular senescence (Figs.1A, B and C). Importantly, upregulated protein synthesis was also observed in replicative senescence in TIG-7 cells (Figs. 1D and E) and in cellular senescence induced by expression of oncogenic Ras in SVts7-1 cells (Figs. 1F and G). These findings indicated that protein synthesis was upregulated during the induction of cellular senescence.

**Figure 1.**
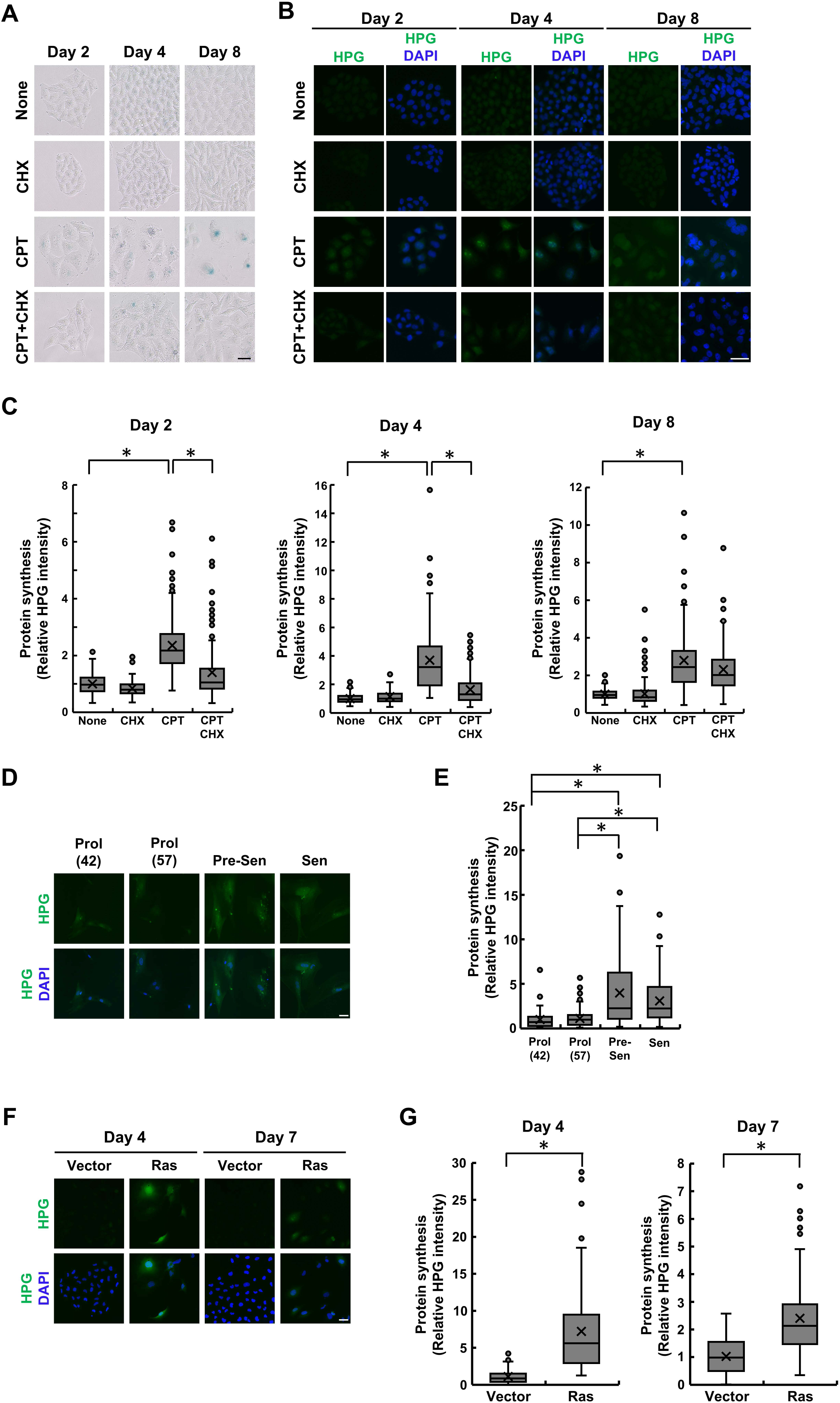
Upregulated protein synthesis during the induction of cellular senescence. A. HeLa cells were cultured with camptothecin (CPT) in the presence or absence of a low dose of cycloheximide (CHX) for the days indicated and were stained with senescence-associated ß-galactosidase (SA-ß-gal). Scale Bars, 50 µm. B, C. The cells in (A) were cultured with L-homopropargylglycine (HPG), a methionine analogue, to label nascent polypeptides. The fluorescence signals of HPG-labelled polypeptides were detected by fluorescence microscopy (B) and quantified (>100 cells) (C). The fluorescence intensity is expressed as a value relative to that of non-treated cells (None). An asterisk indicates statistical significance, *P*<0.05. Scale Bars, 50 µm. D, E. Proliferating (Prol, 42 and 57 PDLs), pre-senescent (Pre-sen, 65 PDLs), or senescent (Sen, 70 PDLs) TIG-7 cells were cultured with HPG to label nascent polypeptides. The fluorescence signals of HPG-labelled polypeptides were detected by fluorescence microscopy (D) and quantified (>100 cells) (E). The fluorescence intensity is expressed as a value relative to that of proliferating cells (42 PDLs). An asterisk indicates statistical significance, *P*<0.05. Scale Bars, 50 µm. PDLs, Population doubling levels. F, G. SVts7-1 cells transfected with a Ras-expressing or an empty vector were cultured with HPG to label nascent polypeptides at the days indicated after transfection. The fluorescence signals of HPG-labelled polypeptides were detected by fluorescence microscopy (F) and quantified (>100 cells) (G). The fluorescence intensity is expressed as a value relative to that of the cells transfected with an empty vector (Vector). An asterisk indicates statistical significance, *P*<0.05. Scale Bars, 50 µm.

### 3.2 Loss of proteostasis in senescent cells

To gain insights into the mechanisms by which protein synthesis regulates cellular senescence, we examined the possibility that upregulated protein synthesis induces a loss of proteostasis, which is indicated by increased protein aggregates that are insoluble in detergent. We found that detergent-insoluble proteins were increased in CPT-induced senescent cells; and that slowdown of protein synthesis by a low dose of cycloheximide suppressed the increase in detergent-insoluble proteins by CPT (Fig. 2A). Importantly, increased detergent-insoluble proteins were also observed in replicative senescent cells (Fig.2B) and in Ras-induced senescent cells (Fig. 2C). Moreover, these results were confirmed by staining aggregated proteins in cells with a fluorescence dye, ProteoStat; that is, increased protein aggregates were observed in CPT-induced senescent cells, replicative senescent cells, and Ras-induced senescent cells (Figs. 2D, E, F, G, H, and I). These results suggested that disturbed proteostasis was induced through upregulated protein synthesis in senescent cells.

**Figure 2.**
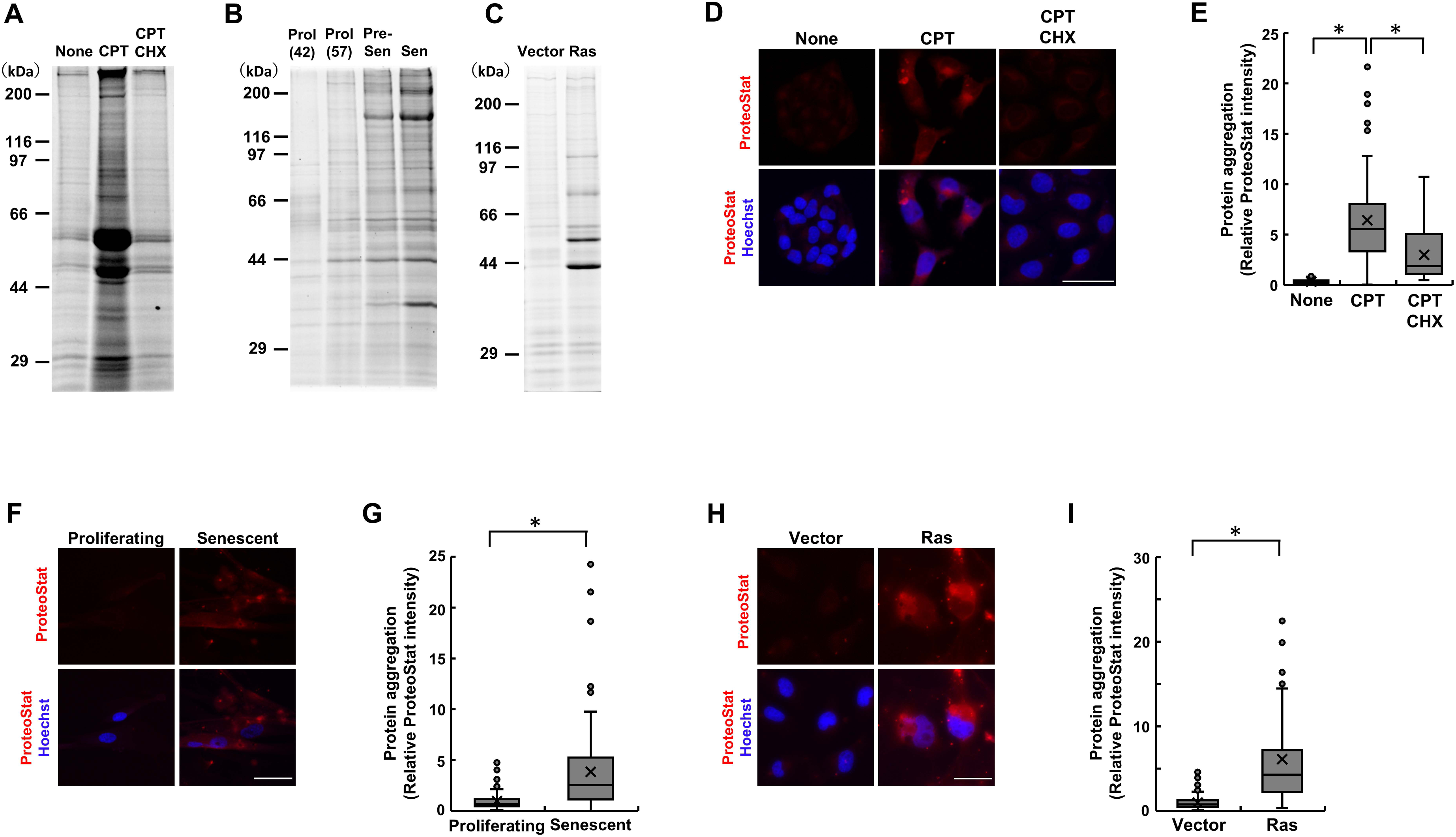
Loss of proteostasis by upregulated protein synthesis. A. Detergent-insoluble proteins were extracted from HeLa cells cultured with camptothecin (CPT) in the presence or absence of a low dose of cycloheximide (CHX) for 6 days and subjected to SDS-PAGE analysis. B. Detergent-insoluble proteins were extracted from proliferating (Prol, 42 and 57 PDLs), pre-senescent (Pre-sen, 65 PDLs), or senescent (Sen, 70 PDLs) TIG-7 cells and subjected to SDS-PAGE analysis. PDLs, Population doubling levels. C. Detergent-insoluble proteins were extracted from the cells transfected with a Ras-expressing or an empty vector 6 days after transfection and subjected to SDS-PAGE analysis. D, E. Protein aggregates were stained with the ProteoStat dye in HeLa cells cultured with CPT in the presence or absence of a low dose of CHX (D) for 6 days and quantified (>100 cells) (E). The fluorescence intensity is expressed as a value relative to that of non-treated cells (None). An asterisk indicates statistical significance, *P*<0.05. Scale Bars, 50 µm. F, G. Protein aggregates were stained with the ProteoStat dye in proliferating or senescent TIG-7 cells (F) and quantified (>100 cells) (G). The fluorescence intensity is expressed as a value relative to that of proliferating cells. An asterisk indicates statistical significance, *P*<0.05. Scale Bars, 50 µm. H, I. Protein aggregates were stained with the ProteoStat dye in the cells transfected with a Ras-expressing or an empty vector (H) 6 days after transfection and quantified (>100 cells) (I). The fluorescence intensity is expressed as a value relative to that of the cells transfected with an empty vector (Vector). An asterisk indicates statistical significance, *P*<0.05. Scale Bars, 50 µm.

### 3.3 Regulation of cellular senescence by a chemical chaperone

The above observations suggested a role of proteostasis in cellular senescence. We then determined whether disturbed proteostasis is causally involved in the induction of cellular senescence. For this, we examined the effect of 4-phenyl butyric acid (4-PBA), a chemical chaperone that promotes refolding of misfolded proteins, on the induction of cellular senescence (Cortez & Sim, 2014). Indeed, 4-PBA decreased protein aggregates in CPT-treated or Ras-expressing cells (Figs. 3A, B, C, and D). Importantly, 4-PBA suppressed the induction of cellular senescence by CPT or Ras, as indicated by increased colony formation and decreased SA-ß-gal positive cells by 4-PBA in CPT-treated or Ras-expressing cells (Figs. 3E, F, G, and H). In addition, 4-PBA significantly extended the replicative lifespan of TIG-7 cells, the result of which indicated the suppression of replicative senescence by 4-PBA (Fig. 3I). These observations suggested that 4-PBA suppressed the induction of cellular senescence through maintaining proteostasis.

**Figure 3.**
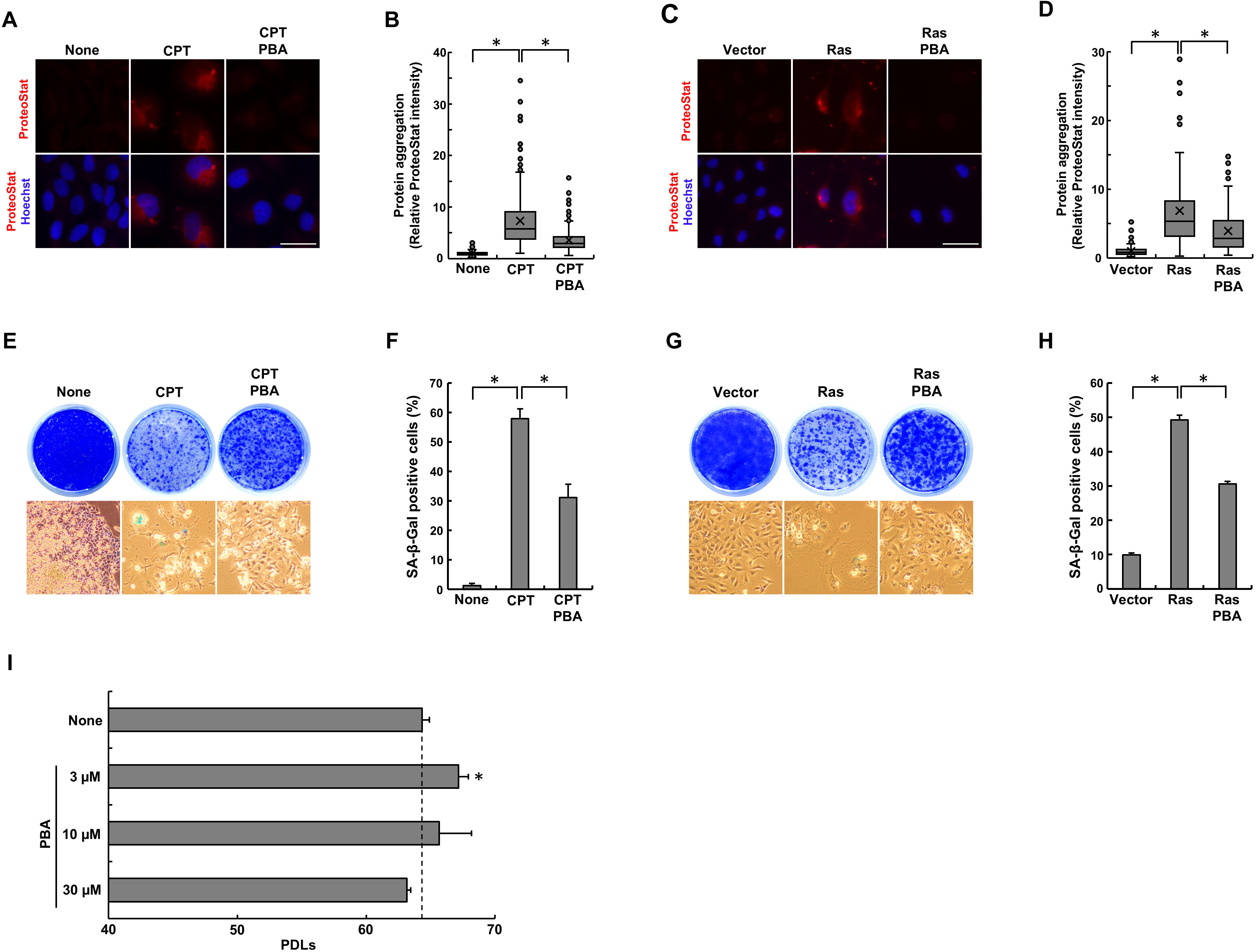
Suppression of cellular senescence by a chemical chaperon. A, B. Protein aggregates were stained with the ProteoStat dye in HeLa cells cultured with camptothecin (CPT) in the presence or absence of 4-PBA for 6 days (A) and quantified (>100 cells) (B). The fluorescence intensity is expressed as a value relative to that of non-treated cells (None). An asterisk indicates statistical significance, *P*<0.05. Scale Bars, 50 µm. C, D. Protein aggregates were stained with the ProteoStat dye in the cells transfected with a Ras-expressing or an empty vector in the presence or absence of 4-PBA 6 days after transfection (C) and quantified (>100 cells) (D). The fluorescence intensity is expressed as a value relative to that of the cells transfected with an empty vector (Vector). An asterisk indicates statistical significance, *P*<0.05. Scale Bars, 50 µm. E, F. HeLa cells were cultured with CPT in the presence or absence of 4-PBA. Cells were stained with SA-ß-gal, and formed colonies were stained with Coomassie Brilliant Blue (CBB) (E). The percentage of the SA-ß-gal-positive cells was determined (>100 cells, n=3) (F). An asterisk indicates statistical significance, *P*<0.05. G, H. Cells transfected with a Ras-expressing or an empty vector were cultured in the presence or absence of 4-PBA. Cells were stained with SA-ß-gal, and formed colonies were stained with CBB (G). The percentage of the SA-ß-gal-positive cells was determined (>100 cells, n=3) (H). An asterisk indicates statistical significance, *P*<0.05. I. TIG-7 cells were cultured with 4-PBA until they ceased to divide. Population doubling levels (PDLs) were determined (n=3). An asterisk indicates statistical significance, *P*<0.05. Scale Bars, 50 µm.

### 3.4 Decreased LBR function by CPT

The above results suggested that cellular senescence was induced at least partly through disturbed proteostasis. We explored the mechanisms of how disturbed proteostasis induces cellular senescence in CPT-treated cells. Interestingly, our previous findings indicated that disturbed proteostasis induces cellular senescence through deceasing the function of LBR, which is a nuclear envelope protein that regulates heterochromatin organization (En, Takauji, Miki, et al., 2020; Olins, Rhodes, Welch, Zwerger, & Olins, 2010). We then examined the implication of LBR in CPT-induced cellular senescence and found that LBR, which is localized at the nuclear envelope in proliferating cells, was decreased at the nuclear envelope in CPT-treated cells (Figs. 4A and B). Importantly, we also found that 4-PBA prevented the decrease of LBR at the nuclear envelope in CPT-treated cells (Figs. 4A and B). These results suggested that CPT decreased LBR function through disturbed proteostasis.

**Figure 4.**
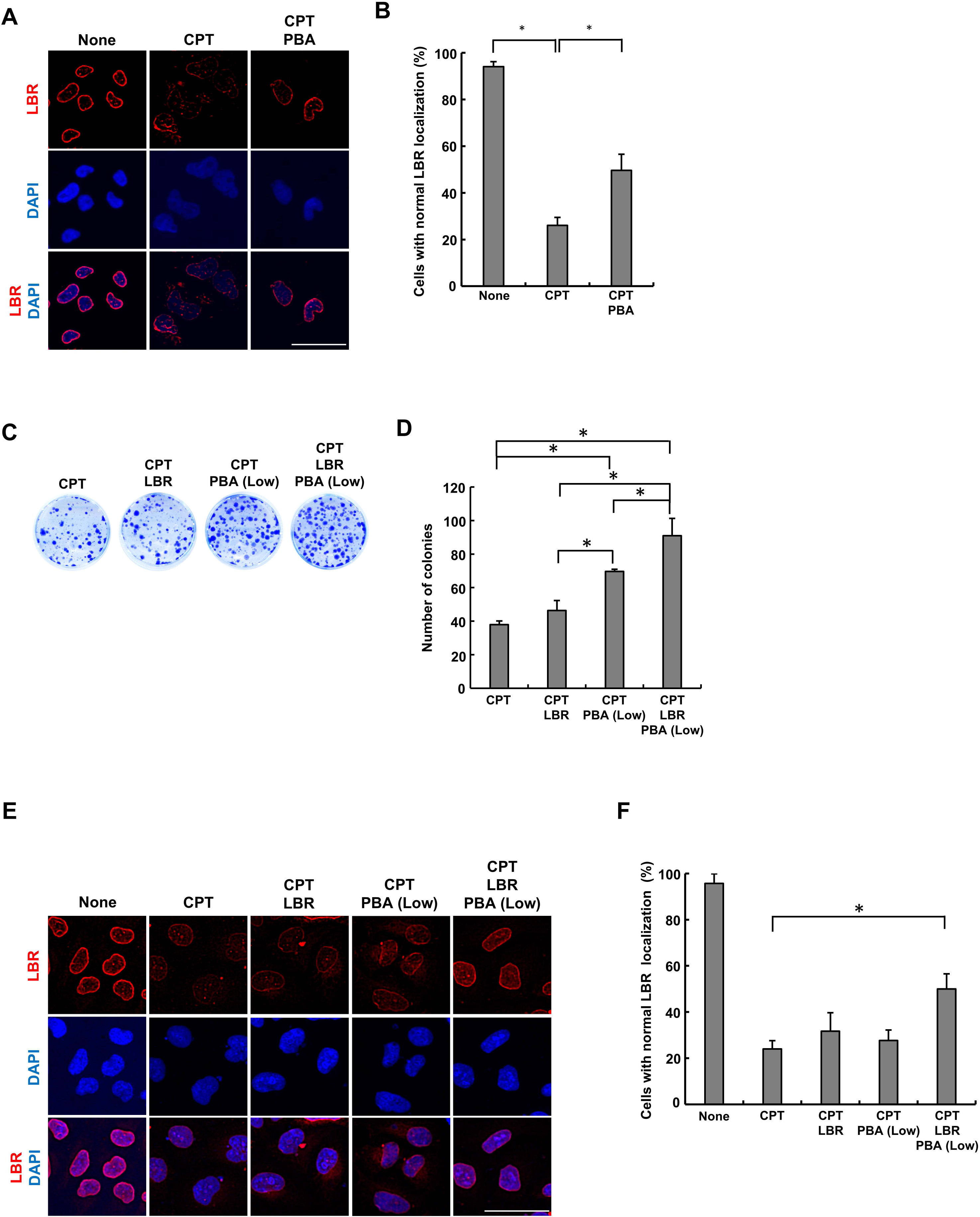
Involvement of LBR in cellular senescence induced by CPT. A, B. Localization of LBR was examined by immunohistochemical analysis with the LBR antibody in HeLa cells cultured with camptothecin (CPT) in the presence or absence of 4-PBA (300 μM) for 6 days (A). The percentage of the cells with normal localization of LBR was determined (>100 cells, n=3) (B). An asterisk indicates statistical significance, *P*<0.05. Scale Bars, 50 µm. C, D. HeLa^T-LBR^ cells were cultured with CPT in the presence or absence of doxycycline (30 ng/ml) or a low dose of 4-PBA (30 μM). Formed colonies were stained with Coomassie Brilliant Blue (C) and counted (n=3) (D). An asterisk indicates statistical significance, *P*<0.05. E, F. Localization of LBR was examined by immunohistochemical analysis with the LBR antibody in HeLa^T-LBR^ cells cultured with CPT in the presence or absence of doxycycline (30 ng/ml) or a low dose of 4-PBA (30 μM) for 6 days (E). The percentage of the cells with normal localization of LBR was determined (>100 cells, n=3) (F). An asterisk indicates statistical significance, *P*<0.05.

### 3.5 Involvement of LBR in cellular senescence induced by CPT

We determined whether decreased LBR function is causally involved in the induction of cellular senescence by CPT. For this, we employed the human cells that increase the expression of *LBR* in a doxycycline-dependent manner, HeLa^T-LBR^ (En, Takauji, Ayusawa, et al., 2020; En, Takauji, Miki, et al., 2020); however, we found that induction of cellular senescence by CPT was barely suppressed by enforced *LBR* expression alone (Figs. 4C and D). We then examined the effect of enforced *LBR* expression on cellular senescence in the presence of a low dose of 4-PBA, which weakly suppressed cellular senescence (Figs. 4C and D). Interestingly, enforced *LBR* expression showed the suppressive effect on CPT-induced cellular senescence in the presence of a low dose of 4-PBA, as indicated by increased colony formation by enforced *LBR* expression in the presence of a low dose of 4-PBA, compared with colony formation in the presence of a low dose of 4-PBA alone (Fig. 4C and D). Consistently, the cells with normal localization of LBR were increased by enforced *LBR* expression in the presence of a low dose of 4-PBA in CPT-treated cells (Figs. 4E and F). LBR, even upon enforced expression, may require the assist of a low dose of 4-PBA to function under proteotoxic stress imposed by CPT. These results indicated that LBR was at least partly involved in the induction of cellular senescence by CPT.

### 3.6 Involvement of LBR in cellular senescence induced by Ras

We additionally examined the involvement of LBR in the induction of cellular senescence by Ras. LBR was decreased at the nuclear envelope in Ras-induced senescent cells as well (Figs. 5A and B), and notably, enforced expression of *LBR* significantly suppressed the induction of cellular senescence by Ras, as indicated by increased colony formation and decreased SA-ß-gal-positive cells by *LBR* expression in Ras-expressing cells (Figs. 5C and D). We also found that the cells with normal localization of LBR were increased by treatment with 4-PBA or *LBR* expression alone (Figs. 5A, B, E and F). These results indicated that decreased LBR function was causally involved in the induction of cellular senescence by Ras as well as by CPT. Suppression of Ras-induced senescence by enforced LBR expression alone without the assist of a low dose of 4-PBA might be due to that Ras induced less severe proteotoxic stress than CPT (Figs. 2A and C).

**Figure 5.**
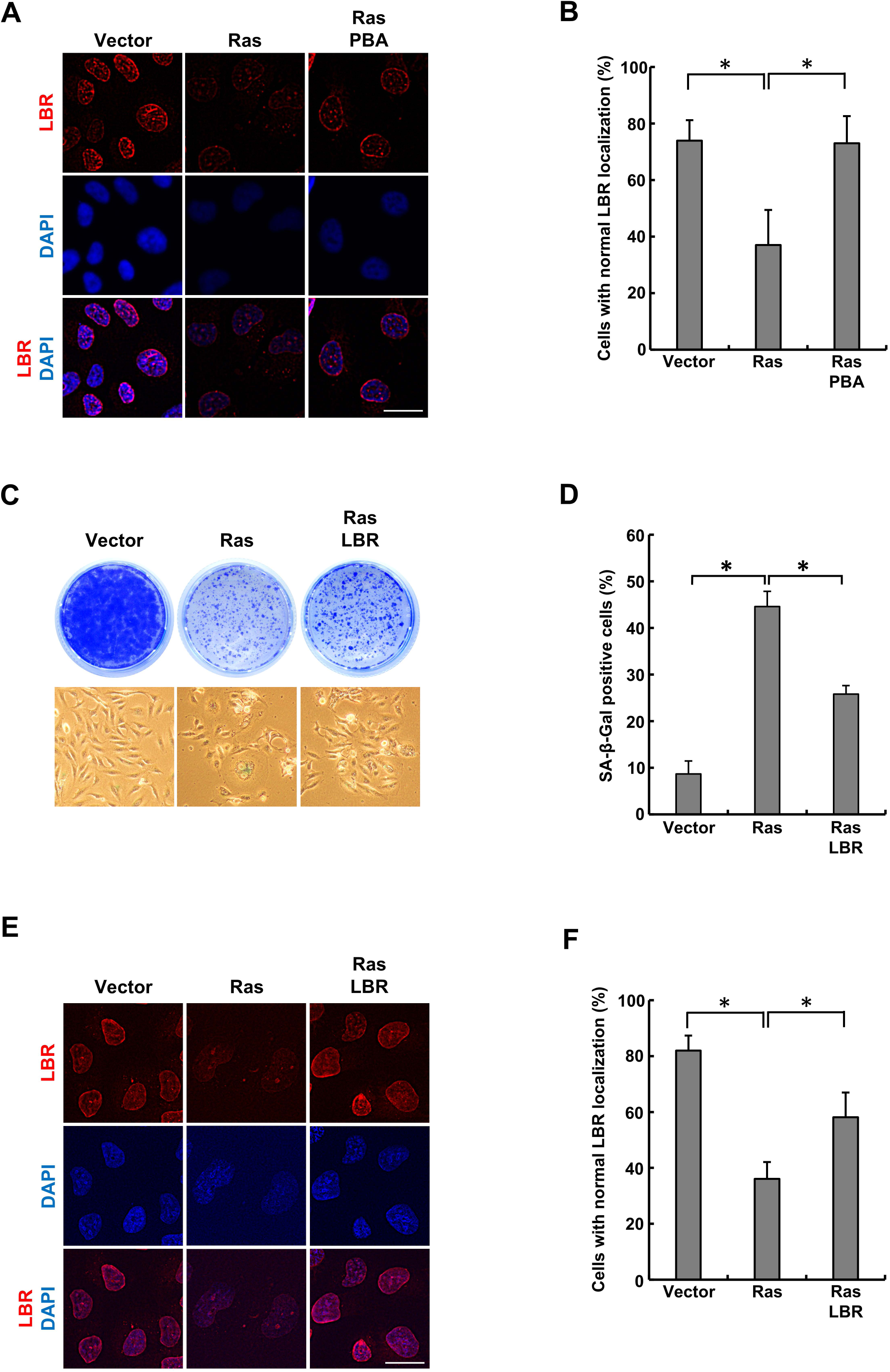
Involvement of LBR in cellular senescence induced by Ras. A, B. Localization of LBR was examined by immunohistochemical analysis with the LBR antibody in Ras-expressing cells in the presence or absence of 4-PBA (A). The percentage of the cells with normal localization of LBR was determined (>100 cells, n=3) (B). An asterisk indicates statistical significance, *P*<0.05. Scale Bars, 50 µm. C, D. Cells were transfected with Ras together with or without *LBR* and cultured. The cells were stained with SA-ß-gal, and formed colonies were stained with Coomassie Brilliant Blue (C). The percentage of the SA-ß-gal positive cells was determined (>100 cells, n=3) (D). An asterisk indicates statistical significance, *P*<0.05. E, F. Localization of LBR was examined by immunohistochemical analysis with the LBR antibody in the cells transfected with Ras together with or without *LBR* (E). The percentage of the cells with normal localization of LBR was determined (>100 cells, n=3) (F). Scale Bars, 50 µm. An asterisk indicates statistical significance, *P*<0.05. Scale Bars, 50 µm.

### 3.7 Altered chromatin configuration and increased DNA damage by decreased LBR function

We examined whether decreased LBR function alters chromatin configuration in cells. For this, cells were treated with DNase I, which more efficiently digests chromatin DNA in open chromatin regions than that in closed chromatin regions. Chromatin DNA was more efficiently digested by DNase I in CPT-treated or Ras-expressing cells than in untreated or empty vector-transfected cells (Figs. 6A, B, C and D). Importantly, we found that 4-PBA suppressed the increased DNase I sensitivity in CPT-treated or Ras-expressing cells (Figs. 6A, B, C and D). Furthermore, enforced expression of *LBR* suppressed the increased DNase I sensitivity in Ras-expressing cells (Figs. 6E and F). Thus, disturbed proteostasis loosened chromatin configuration through decreased LBR function.

**Figure 6.**
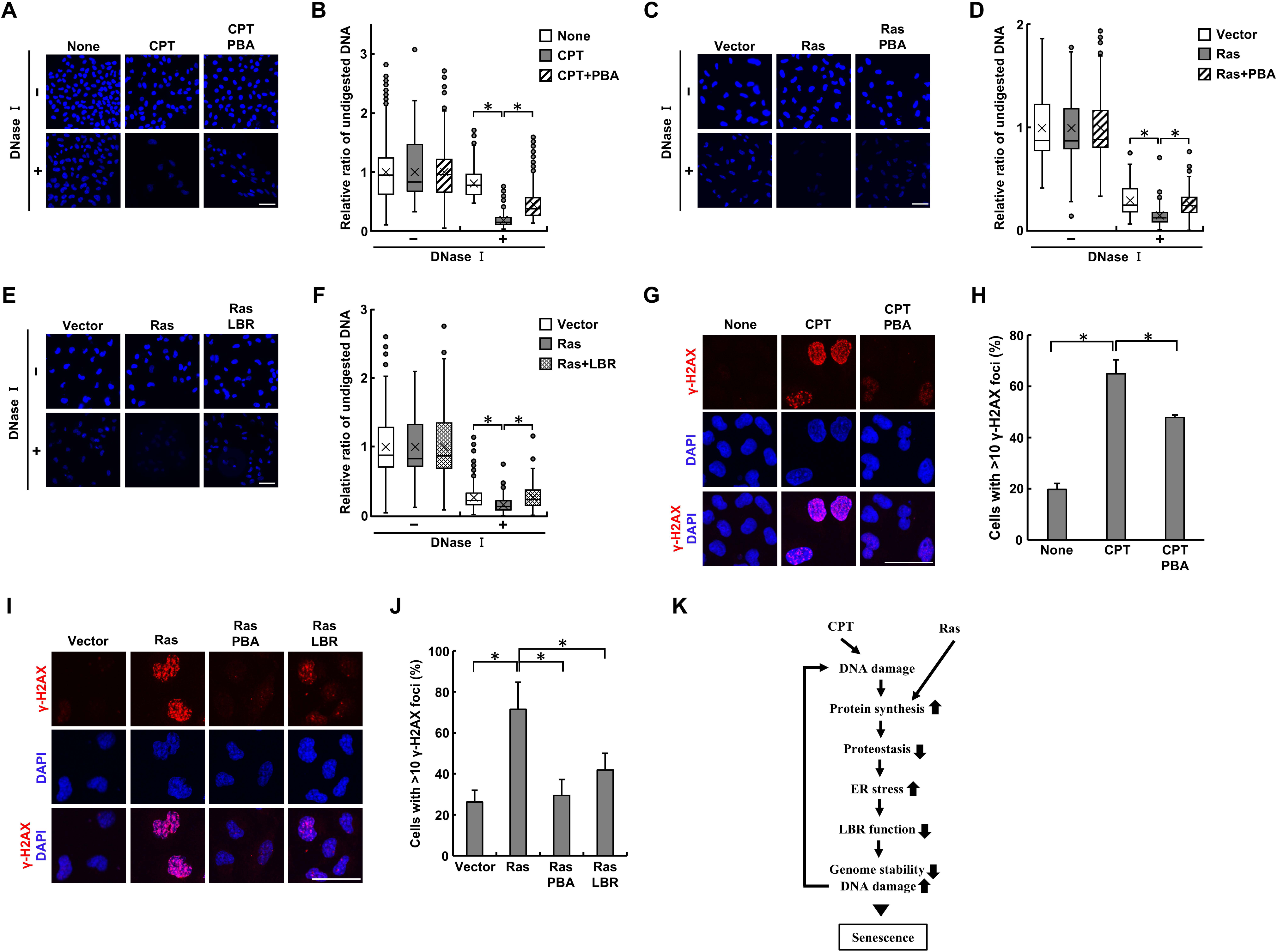
Altered chromatin configuration and increased DNA damage by decreased LBR function. A, B. HeLa cells were cultured with camptothecin (CPT) in the presence or absence of 4-PBA and treated with DNase I. Residual DNA was stained with DAPI (A) and quantified (>100 cells) (B). The amount of residual DNA after DNase I treatment is expressed as a value relative to that of DNA without DNase I treatment. An asterisk indicates statistical significance, *P*<0.05. Scale Bars, 50 µm. C, D. Cells transfected with Ras were cultured in the presence or absence of 4-PBA and treated with DNase I. Residual DNA was stained with DAPI (C) and quantified (>100 cells) (D). The amount of residual DNA after DNase I treatment is expressed as a value relative to that of DNA without DNase I treatment. An asterisk indicates statistical significance, *P*<0.05. Scale Bars, 50 µm. E, F. Cells transfected with Ras together with or without *LBR* were treated with DNase I. Residual DNA was stained with DAPI (C) and quantified (>100 cells) (D). The amount of residual DNA after DNase I treatment is expressed as a value relative to that of DNA without DNase I treatment. An asterisk indicates statistical significance, *P*<0.05. Scale Bars, 50 µm. G, H. DNA damage was detected by immunohistochemical analysis with the γ-H2AX antibody in HeLa cells cultured with CPT in the presence or absence of 4-PBA (E). The percentage of the cells with >10 γ-H2AX foci/nuclei was determined (>100 cells) (F). An asterisk indicates statistical significance, *P*<0.05. Scale Bars, 50 µm. I, J. DNA damage was detected by immunohistochemical analysis with the γ-H2AX antibody in the cells transfected with Ras together with or without *LBR* in the presence or absence of 4-PBA (G). The percentage of the cells with >10 γ-H2AX foci/nuclei was determined (>100 cells) (H). An asterisk indicates statistical significance, *P*<0.05. Scale Bars, 50 µm. K. Schematic representation of the mechanisms of cellular senescence is shown.

We then examined the mechanisms of how decreased LBR function induces cellular senescence in CPT-treated or Ras-expressing cells. CPT induced the formation of γ-H2AX foci, which represents DNA damage (Figs. 6G and H). This observation is reasonable because CPT is a reagent that generates DNA strand breaks; however, interestingly, increased γ-H2AX foci formation by CPT was significantly suppressed by simultaneous treatment with 4-PBA (Figs. 6G and H). Similarly, γ-H2AX foci formation was increased by Ras, and its increase was significantly suppressed by simultaneous treatment with 4-PBA (Figs. 6I and J). Furthermore, enforced *LBR* expression significantly suppressed the increased γ-H2AX foci formation by Ras (Figs. 6I and J). These results suggested that disturbed proteostasis induced DNA damage through decreased LBR function.

Collectively, these findings indicated that disturbed proteostasis induced cellular senescence through decreased LBR function that altered chromatin configuration and increased DNA damage. Given the role of DNA damage in cellular senescence, increased DNA damage by decreased LBR function was at least partly causative for the induction of cellular senescence.

### 3.8 ER stress upon induction of cellular senescence

We investigated the mechanism for decreased LBR function by CPT or Ras. Our previous study showed that disturbed proteostasis in endoplasmic reticulum (ER), *i.e.*, ER stress, decreases LBR function, consistent with the fact that LBR is a nuclear envelope protein that is transferred from ER to the nuclear envelope after being properly folded in ER (Ellenberg et al., 1997; En, Takauji, Miki, et al., 2020). We then examined the expression of the genes involved in ER stress response and found that ER stress was induced by CPT or Ras (Supplementary fig. 1). This result suggested that disturbed proteostasis in ER may cause the decreased LBR function and suggested the key role of proteostasis in ER in cellular senescence.

## 4 Discussion

In the present study, we aimed to reveal the conserved mechanisms underlying cellular senescence. We have previously shown that a low dose of cycloheximide, which slows down protein synthesis, effectively suppressed cellular senescence induced by various types of stress (En, Takauji, Miki, et al., 2020; Sumikawa et al., 2005; Takauji, En, et al., 2016; Takauji, Wada, et al., 2016). In the present study, we further showed that protein synthesis was upregulated during the induction of cellular senescence. These results suggested the crucial roles of protein synthesis in the regulation of cellular senescence. Of note, these results appear to be in agreement with the reports that cellular senescence is suppressed by decreased activity of mTOR, a master regulator of protein synthesis; and that mTOR is activated at the onset of the induction of cellular senescence (Cho & Hwang, 2012; Herranz et al., 2015; Leontieva & Blagosklonny, 2010; Narita et al., 2011). Interestingly, upregulated protein synthesis is also reported in cells from progeria syndrome patients, compared with those from healthy controls (Buchwalter & Hetzer, 2017). These studies indicate the important roles of upregulated protein synthesis in the induction of cellular senescence. Additionally, it is worthwhile mentioning that organismal aging is suppressed by decreased protein synthesis due to defects in the proteins such as mTOR, ribosomal protein S6 kinase, translation initiation factors eIF4e/f/g, or ribosomal proteins in yeast, *C. elegans*, *Drosophila melanogaster*, or mice (Demidenko et al., 2009; Harrison et al., 2009; Kaeberlein et al., 2005; Kapahi et al., 2004; Pan et al., 2007; Steffen et al., 2008; Syntichaki, Troulinaki, & Tavernarakis, 2007; Vellai et al., 2003). Thereby, protein synthesis regulates organismal aging as well as cellular senescence. However, inconsistent results have been reported on protein synthesis in senescent cells: some studies report the upregulation of protein synthesis in senescent cells and others report the downregulation of it (Herranz et al., 2015; Lee et al., 2021; Srikantan, Marasa, Becker, Gorospe, & Abdelmohsen, 2011; Wu et al., 2019). This may be due to that cells undergoing cellular senescence may show upregulated protein synthesis but those that have entered deep senescence may show downregulated protein synthesis.

How dose upregulated protein synthesis induce cellular senescence? Upregulated protein synthesis consumes a large amount of ATP and consequently leads to mitochondrial exhaustion with increased reactive oxygen species (ROS) production which accelerates cellular senescence. Indeed, our previous study indicated that senescent cells showed increased ROS production, and a low dose of cycloheximide prevented it (Takauji, En, et al., 2016). However, the study also indicated that prevention of the increased ROS production is not a major cause for the suppressive effect of cycloheximide on the induction of cellular senescence (Takauji, En, et al., 2016). Here we showed that upregulated protein synthesis induced disturbed proteostasis that played important roles in the induction of cellular senescence. Notably, this is consistent with many lines of evidence that proteostasis is lost in senescent cells or aged organisms (David et al., 2010; Sabath et al., 2020; Tanase et al., 2016; Walther et al., 2015).

It seems to be reasonable that disturbed proteostasis induces cellular senescence, given that a variety of proteins are inactivated by proteotoxic stress. However, it is not clearly determined which protein inactivation is crucial for the induction of cellular senescence by disturbed proteostasis. We have previously shown that disturbed proteostasis induced cellular senescence by decreasing the function of LBR, a nuclear envelope protein that regulates heterochromatin organization (En, Takauji, Miki, et al., 2020; Olins et al., 2010). Consistently, we here showed that LBR function was decreased by CPT or Ras, and decreased LBR function was causally involved in the induction of cellular senescence by them. Thus, LBR is one of the important proteins that are inactivated by disturbed proteostasis and induce cellular senescence upon inactivation. Since ER stress causes aberrant localization of LBR (En, Takauji, Miki, et al., 2020), it is likely that disturbed proteostasis in ER (*i.e.*, ER stress) interfered with the proper folding of LBR and consequently caused decreased LBR function. This view is consistent with the fact that LBR is a nuclear envelope protein that is transferred from ER to the nuclear envelope after being properly folded in ER (Ellenberg et al., 1997). We further explored the mechanism by which decreased LBR function induces cellular senescence and showed that decreased LBR function altered chromatin configuration and increased DNA damage. Given the role of DNA damage in cellular senescence, increased DNA damage was at least partly involved in the induction of cellular senescence by disturbed proteostasis. Thereby, these findings suggest that disturbed proteostasis induced cellular senescence through decreased LBR function that altered chromatin configuration and increased DNA damage. This is consistent with our previous finding that LBR regulates genome stability by modulating chromatin configuration (En et al., 2024).

We summarized our findings in Figure 6K. Our findings indicated that i) cells undergoing cellular senescence showed upregulated protein synthesis that led to disturbed proteostasis; ii) disturbed proteostasis decreased the function of LBR; and iii) decreased LBR function caused altered chromatin configuration and increased DNA damage, which leads to cellular senescence. It is interesting to note that DNA damage was increased in a positive feedback loop manner and reinforced the induction of cellular senescence. Our findings revealed a molecular link between protein synthesis and chromatin organization and accounted for the phenotypes of senescent cells which show disturbed proteostasis, altered chromatin organization, and increased genome instability. Our findings provided the general model for the mechanisms of cellular senescence.

## Abbreviations

SA-ß-gal: senescence-associated ß-galactosidase
SASP: senescence-associated secretory phenotype
CPT: camptothecin
4-PBA: 4-phenyl butyric acid

## Author Contributions

YT, KT, HM, TK, SF, and AE conducted the experiments, and MF supervised the project.

## Conflicts of Interest

The authors declare no conflict of interest.

## Acknowledgements

We thank Prof. Ide, T. for providing us with SVts7-1 cells. This work was supported in part by grants-in-aid for Scientific Research from the Ministry of Education, Culture, Sports, Science, and Technology of Japan (20K19643 and 22K11730).

## 6 Figure legends

**Supplementary Fig. 1.**
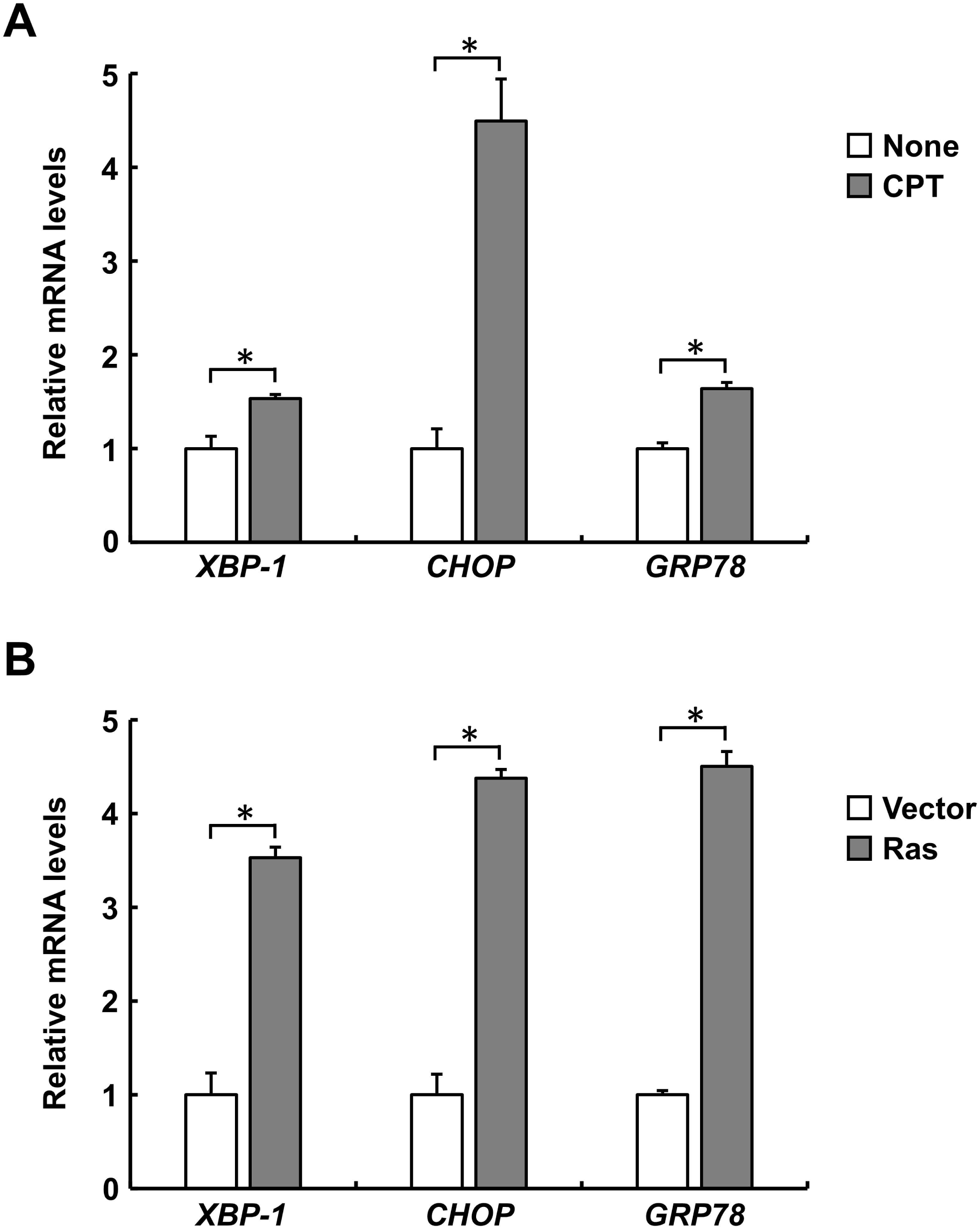
ER stress in cells treated with CPT or Ras. Expression of the genes involved in ER stress response was determined by quantitative RT-PCR in the cells treated with camptothecin (CPT) or transfected with Ras. An asterisk indicates statistical significance, *P*<0.05.

## Notes

### Competing Interest Statement

The authors have declared no competing interest.

